# SNAP MRI Reveals Association Between Distal Cerebral Arterial Flow and Cognitive Function in an Aging Population

**DOI:** 10.64898/2026.03.21.713392

**Authors:** Xiaodong Ma, Vincent Koppelmans, Halit Akcicek, Ebru Yaman Akcicek, Jincheng Shen, Li Chen, Niranjan Balu, Chun Yuan, Jace B. King

## Abstract

**Objective:** Impaired blood flow has recently been recognized as a critical contributor to cognitive impairment and dementia. It was reported that cerebral distal arterial flow measured from Simultaneous Non-contrast Angiography and Intraplaque Hemorrhage (SNAP) MRI is associated with post-treatment cognitive function improvement in carotid atherosclerosis patients. In this study, we aim to evaluate the value of SNAP-based measurements in assessing cerebrovascular function in an aging population.

**Materials and Methods:** Neurovascular MRI data were collected on 36 aging participants (22 cognitively unimpaired and 14 impaired; 9 mild cognitive impairment (MCI) and 5 Alzheimer’s Disease (AD)). Neurovascular MRI measurements, including white matter hyperintensities (WMH) volumes, cerebral blood flow (CBF), and SNAP-based distal cerebral arterial flow (dCAF) index, were quantified. Cognitive function was assessed using the Repeatable Battery for the Assessment of Neuropsychological Status (RBANS).

**Results:** Significant differences in the dCAF index were observed between cognitively unimpaired and impaired groups, and the dCAF index was significantly correlated with the RBANS total score. While CBF was significantly associated with dCAF index, there is no significant correlation of CBF or WMH with the RBANS score in this population.

**Conclusion:** Our findings suggest that the dCAF measured with SNAP MRI is valuable for evaluating the cognition-related cerebrovascular condition in an aging population.

## 1 Introduction

Alzheimer’s disease (AD) and related dementias and cerebrovascular diseases share common risk factors[1–4] and frequently co-occur, as evidenced by autopsy studies[5–10] and clinical diagnoses[11]. Investigating the role of cerebrovascular health in cognitive impairment is critical for understanding vascular contributions to cognitive impairment and dementia (VCID), potentially facilitating improved diagnosis, evaluation, and treatment of dementia.

Decreased blood flow to brain tissue is typically considered a crucial factor linking cerebrovascular disease to cognitive impairment. It has been reported that AD patients have reduced cerebral blood flow (CBF), a.k.a. hypoperfusion, compared to typically aging adults, though the causal relationship between hypoperfusion and AD is still under debate[12–15]. Besides CBF, cerebral arterial flow has drawn increasing attention since it may directly capture blood flow changes due to vessel lesions (such as atherosclerosis), and can therefore assess additional or earlier effects of cerebrovascular factors related to cognitive impairments and AD[16, 17].

Recently, the Simultaneous Non-contrast Angiography and Intraplaque Hemorrhage (SNAP) MRI technique was applied to measure distal cerebral arterial flow (dCAF). SNAP was originally developed as a hyperintensity T1-weighted sequence for improved diagnosis of intra-plaque-hemorrhage, and was recently found sensitive to the slow blood traveling time from proximal arteries to the distal arteries[18, 19]. The underlying principle is that, in SNAP, if blood travels slowly from proximal to distal arteries, a portion of the distal arteries will become invisible[18]. As a result, the total artery length of visible arteries in SNAP can represent dCAF in the brain, and it has been shown to correlate with cognitive outcome in carotid atherosclerotic patients[18–20]. Compared with other conventional cerebral arterial flow measurement techniques, such as 4D flow MRI or dynamic MR angiography, it has several advantages, including efficient acquisition and high spatial resolution. Nonetheless, studies using this measurement have only been conducted in patients with carotid atherosclerosis, and has not yet been applied in an aging population or in AD patients.

In this study, we aim to evaluate distal cerebral arterial flow (dCAF) in an aging population and to investigate its relationship with brain cognition. Specifically, we conducted a cross-sectional MRI study on older participants (aged 65 or older) who were cognitively unimpaired or cognitively impaired (mild cognitive impairment (MCI) or AD). For each participant, neurovascular markers including dCAF, CBF, and white matter hyperintensity (WMH) volume – an established cerebral small vessel disease marker[21, 22] – were measured from the MRI data. In addition, cognitive function was assessed using the Repeatable Battery for the Assessment of Neuropsychological Status (RBANS), which is a brief neuropsychological screening measure assessing language, attention, visuospatial/constructional faculties, and immediate and delayed memory[23]. Extensive research has validated use of the RBANS in cognitively unimpaired[24–27], and cognitively impaired (MCI and AD) older adults[28–30], and was recently shown to correlate with resting-state functional connectivity strength between the default mode and central executive networks (cognitively unimpaired, MCI, and AD) (King, submitted). We then performed group comparisons and correlational analyses among those neurovascular markers and RBANS scores.

## 2 Materials and Methods

### 2.1 Subjects

This research was approved by the Institutional Review Board at the University of Utah. Participants were recruited based on prior involvement in an associated study[31, 32]. All participants provided written informed assent or consent before taking part in the study. Eligibility required participants to be 65 years or older and to have a knowledgeable collateral source who could discuss their cognitive abilities and daily functioning.

Thirty-six older adults (22 cognitively unimpaired; 14 cognitively impaired (5 with AD and 9 with MCI). were enrolled in the current study, which is a subset of participants derived from a larger neuroimaging study who were willing to undergo additional neurovascular MRI scans. Exclusion criteria included: history of major stroke, head injury or other neurological disorder or systemic illness that would likely affect cognition; current or past major psychiatric illness; history of substance abuse; current use of antipsychotics or anticonvulsant medications; and currently residing in a nursing home or other skilled nursing facility, or a history of hallucinations or tremor to exclude potential MCI due to Lewy bodies.

### 2.2 Group Assignment and Cognitive Evaluation

Group assignment was determined at the time of inclusion in an associated study[31, 32], which in turn was based on ADNI protocols[33]. In brief, the Alzheimer’s Disease Neuroimaging Initiative (ADNI2) assessment battery was used to classify participants into cognitively intact, MCI, or AD groups. Due to the limited sample size, we combined MCI and AD participants into a single cognitively impaired sample.

Neuropsychological status was evaluated on average 26.6 +/- 29.3 (min = 0, max = 123) days before MRI, using the Repeatable Battery for the Assessment of Neuropsychological Status (RBANS)[23]. The RBANS evaluates five cognitive domains (immediate memory, visuospatial constructional abilities, language, attention, and delayed memory). RBANS Index scores are based on age-adjusted normative data, yielding standard scores with a mean of 100 and a standard deviation of 15, with higher scores indicating better cognition. All RBANS Index and Total Scale scores were included in the analysis.

### 2.3 MRI Data Acquisition

MRI data were acquired on a Siemens 3T MRI scanner (Prisma-fit, Erlangen, Germany) using a 64-channel head-neck coil. The imaging protocol includes four MRI sequences: 1) Magnetization Prepared Rapid Gradient Echo (MPRAGE), 2) T2-weighted Fluid Attenuated Inversion Recovery (T2-FLAIR), 3) 2D Pseudo-Continuous Arterial Spin Labeling (pCASL), and 4) SNAP. Detailed imaging parameters are shown in Table 1.

**Table 1.** Imaging parameters for MRI sequences.

| equences | Acq./Interp. Voxel Size<br>(mm<sup>3</sup>) | FOV (mm<sup>3</sup>) | TE/TR/TI (ms) | Flip<br>Angle (°) | Accele-<br>ration | Bandwidth<br>(Hz/pixel) | Scan Time<br>(min:sec) | Number of<br>Measurements |
| --- | --- | --- | --- | --- | --- | --- | --- | --- |
| IPRAGE | 0.8×0.8×0.8/- | 256×240×208 | 2.2/2500/1000 | 8 | iPAT=2 | 220 | 6:51 | 1 |
| 2 FLAIR | 1×1×1.25/0.5×0.5×1 | 250×250×176 | 394/6000/1850 | Variable | iPAT=3,<br>PF=6/8 | 781 | 3:26 | 1 |
| D pCASL | 3.0×3.0×5.0/- | 240×240×125 | 11/5300/- | 90 | iPAT=2,<br>PF=6/8 | 2404 | 4:45 | 51 |
| NAP | 0.8×0.8×0.8/0.4×0.4×0.4 | 180×180×70 | 5.7/1830/500 | 11, 5 | - | 399 | 6:23 | 1 |

### 2.4 Image Processing and Quantification

#### 1) White Matter Hyperintensities (WMH)

White matter hyperintensities (WMH) regions were automatically segmented using Samseg in FreeSurfer[34] (Figure 1A), from the combination of MPRAGE and T2-FLAIR images. The WMH regions were further classified into periventricular and deep WMH (i.e., pWMH and dWMH) based on a 10-mm distance rule[35], generating total, pWMH and dWMH volumes by calculating the number of voxels. These WMH volumes were normalized by intracranial volume, also obtained using Samseg in FreeSurfer, following the literature[36]{Cerri, 2023 #1735}.

**Figure 1.**
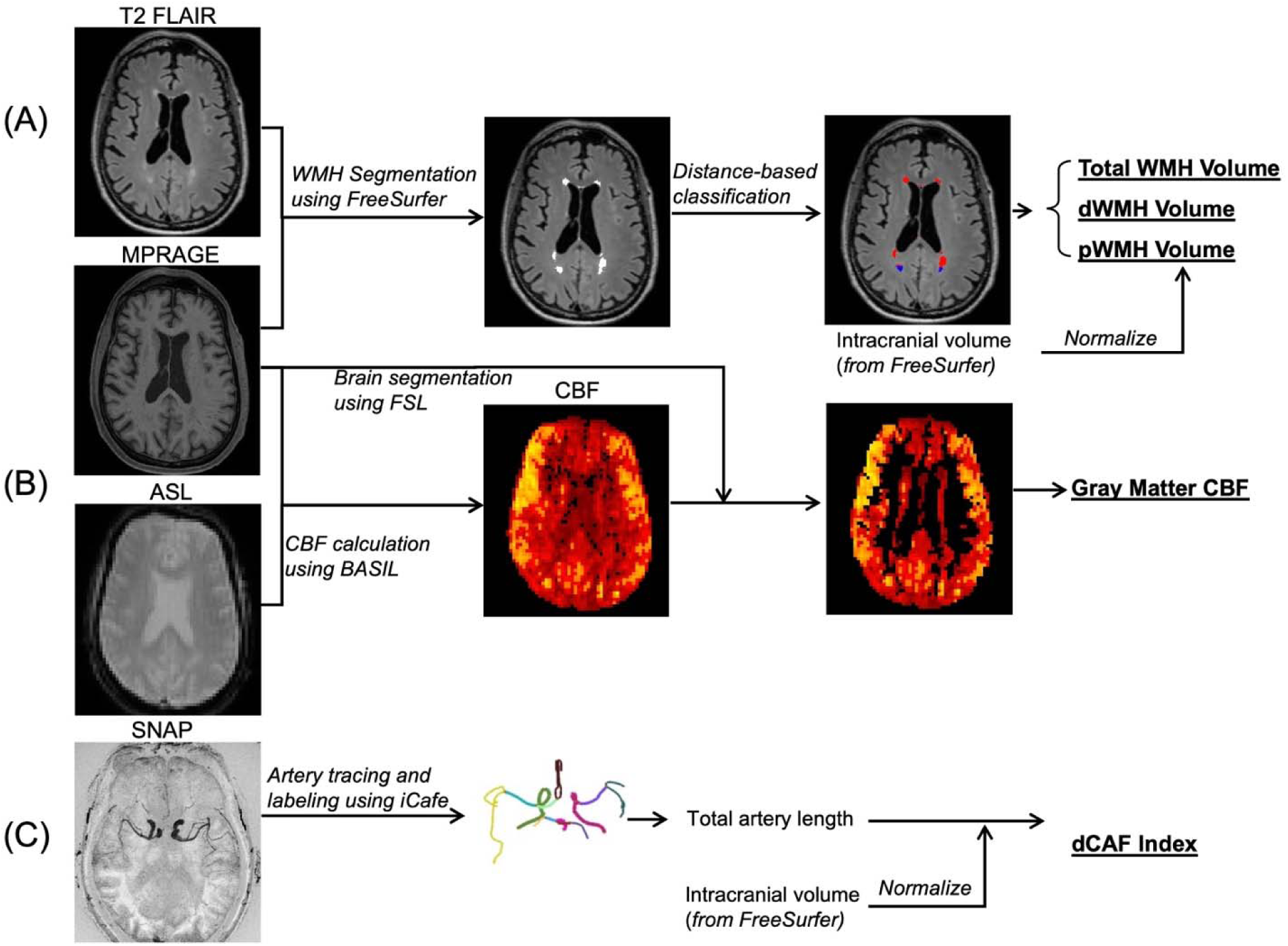
Image processing pipeline for measuring neurovascular markers from MRI data. (A) White matter hyperintensity (WMH) volume (normalized by intracranial volume), (B) gray matter cerebral blood flow (CBF), (C) distal cerebral arterial flow (dCAF) index – defined as the total visible distal artery length. Gray matter CBF and dCAF index were measured in the whole brain as well as in each arterial territory. Arteries were manually traced and labeled on SNAP images using iCafe.

#### 2) Cerebral Blood Flow (CBF)

The pCASL data were processed using BASIL[37] (Figure 1B), with the MPRAGE being used as the structural reference. Motion correction and partial volume correction were applied following the consensus recommendation[38]. A gray matter CBF map was generated for each participant, as it is more reliable compared to white matter CBF[39]. The whole brain gray matter CBF values were computed as the mean value within the gray matter mask generated from MPRAGE. The CBF map was registered to a brain arterial territories atlas[40], and CBF value in each arterial territory including the left or right anterior communicating artery (ACA), middle cerebral artery (MCA), and posterior cerebral artery (PCA) was computed.

#### 3) Distal Cerebral Arterial Flow (dCAF) Index

SNAP data were analyzed by a trained physician reviewer and verified by another physician reviewer using iCafe[41] (Figure 1C), with all cerebral arteries visible in SNAP being manually traced and labeled. Distal arterial length was quantified by summing the lengths of all visible distal arteries (defined as M2, M3, A2, and P2 arteries) in the whole brain. To adjust for brain size, the total arterial length was normalized using intracranial volume (measured by Samseg in FreeSurfer as mentioned above), leading to a normalized total arterial length, called the distal cerebral arterial flow (dCAF) index in this study. With labeled arteries, the dCAF index in each arterial territory (left or right ACA, MCA, and PCA) can also be obtained, computed as territorial distal artery length normalized by intracranial volume.

### 2.5 Statistical Analysis

The Student’s t-test was employed to compare the MRI measurements (WMH volumes, GM-CBF, and dCAF index) between the cognitively unimpaired and impaired groups.

Pearson correlations were used to assess the associations among neurovascular MRI measurements and RBANS Index and Total scores (Model 1). No significant correlations were found between age, sex, and any MRI measurements; therefore, these variables were not included in the model. However, because of the modest sample size and the limited understanding of the relationship between age, sex, and SNAP variables, a post hoc analysis was performed to explore the effects of age and sex (Model 2). The statistical results were considered significant if the p-value was <0.05. All statistical analyses were conducted in MATLAB (The MathWorks, Inc., 2023b) using complete case analysis, excluding observations with missing values (1 missing value for each measurement).

## 3 Results

### 3.1 Summary of Demographic Information and Cognitive Outcome

Demographic information and RBANS scores for all participants are summarized in Table 2. In brief, 36 older participants were included, with 20 females and 16 males, ranging in age from 67.6 to 86.7 (mean age: 77.8). Twenty-two participants were cognitively unimpaired and 14 were cognitively impaired (i.e., diagnosed with MCI or AD; 5 AD and 9 MCI). One cognitively impaired participant was missing an RBANS score and was therefore excluded from the statistical analysis.

**Table 2.**
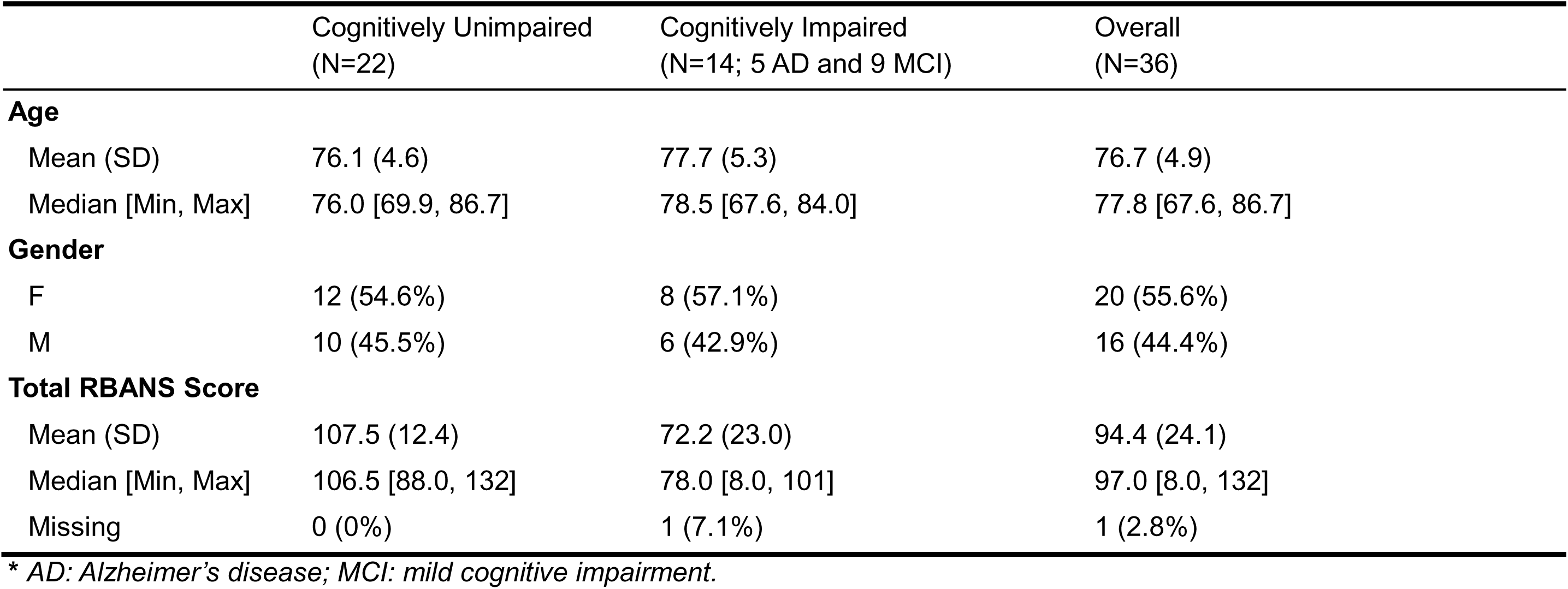
Summary of Demographic and Repeated Battery for the Assessment of Neuropsychological Status Scores.

|  | Cognitively Unimpaired<br>(N=22) | Cognitively Impaired<br>(N=14; 5 AD and 9 MCI) | Overall<br>(N=36) |
| --- | --- | --- | --- |
| **Age** |  |  |  |
| Mean (SD) | 76.1 (4.6) | 77.7 (5.3) | 76.7 (4.9) |
| Median [Min, Max] | 76.0 [69.9, 86.7] | 78.5 [67.6, 84.0] | 77.8 [67.6, 86.7] |
| **Gender** |  |  |  |
| F | 12 (54.6%) | 8 (57.1%) | 20 (55.6%) |
| M | 10 (45.5%) | 6 (42.9%) | 16 (44.4%) |
| **Total RBANS Score** |  |  |  |
| Mean (SD) | 107.5 (12.4) | 72.2 (23.0) | 94.4 (24.1) |
| Median [Min, Max] | 106.5 [88.0, 132] | 78.0 [8.0, 101] | 97.0 [8.0, 132] |
| Missing | 0 (0%) | 1 (7.1%) | 1 (2.8%) |

\* AD: Alzheimer's disease; MCI: mild cognitive impairment.

### 3.2 Groupwise Comparison of Neurovascular MRI Measurements

Compared with the cognitively unimpaired group, the impaired group had a significantly smaller whole-brain dCAF index (*p*=0.004). No significant differences in other whole-brain neurovascular measurements (including pWMH, dWMH, total WMH volumes, and gray matter CBF) were found between the two groups (Figure 2). In addition, the cognitively impaired group had a significantly smaller dCAF index in all arterial territories except left PCA; no significant differences in territorial gray matter CBF were found between the two groups (Figure S1).

**Figure 2.**
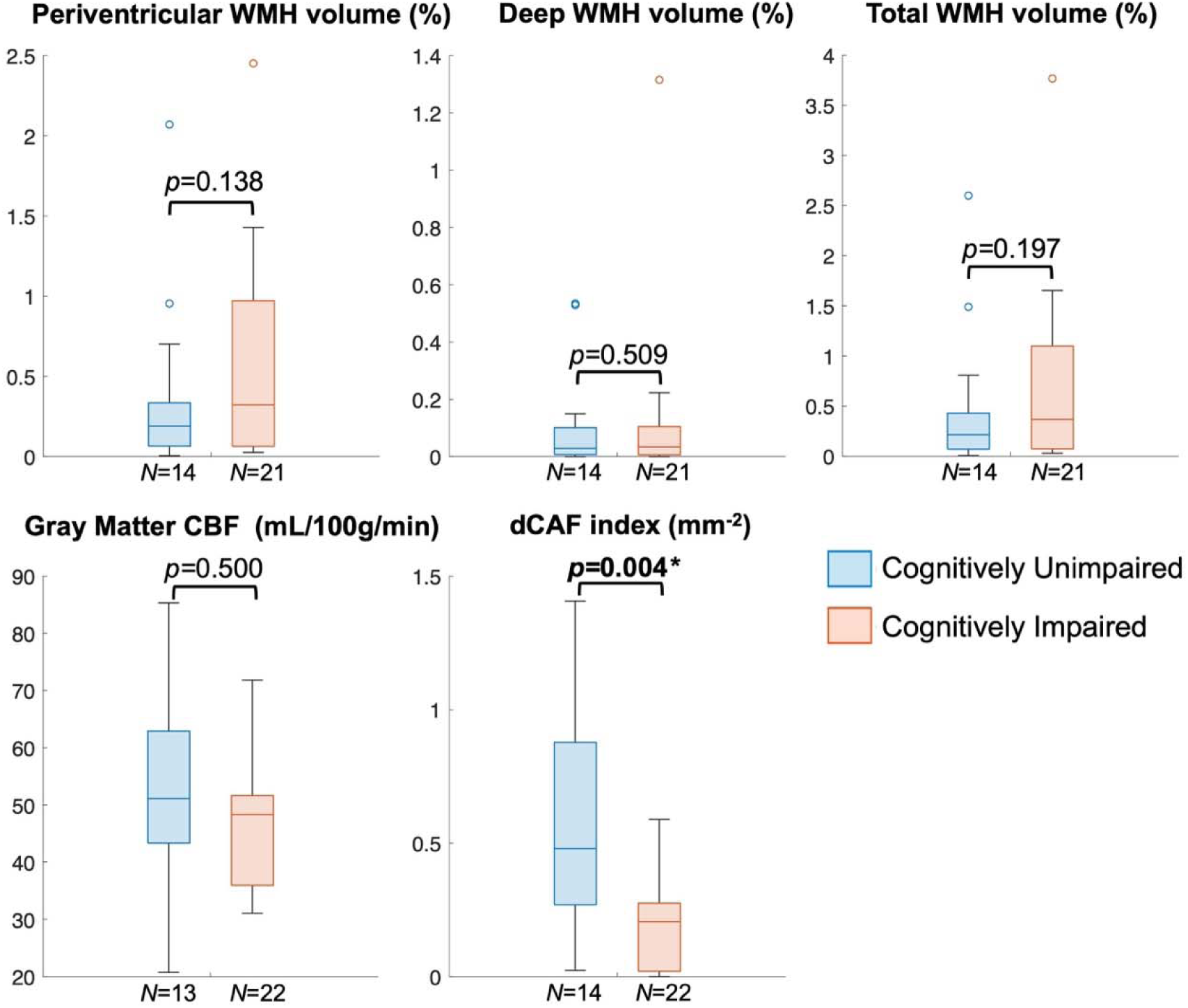
Comparison of whole-brain cerebrovascular MRI measurements between cognitively unimpaired and impaired groups. Shown is the box plot of each measurement displaying the median, upper and lower quartiles. There is a significant difference in the distal cerebral arterial flow (dCAF) index between the two groups.

The difference in dCAF index between cognitively unimpaired and impaired participants is directly visualized in Figure 3, which displays the original SNAP image and the artery tracing and labeling results for two representative participants (one cognitively unimpaired and one cognitively impaired). The cognitively impaired participant has fewer and shorter visible distal cerebral arteries due to a longer blood transition time.

**Figure 3.**
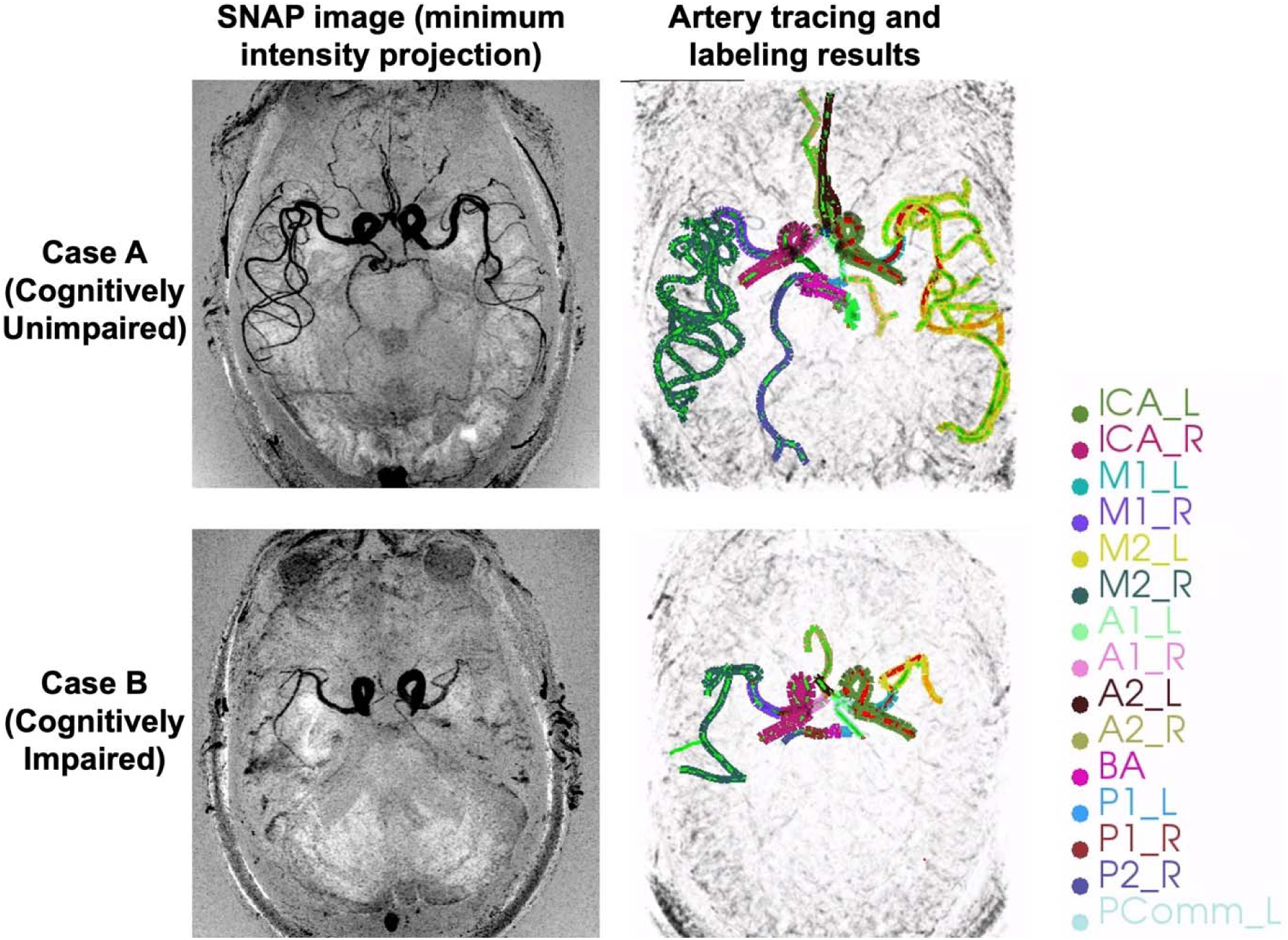
The original SNAP image and the artery tracing and labeling results of one representative cognitively unimpaired participant and one cognitively impaired participant. The cognitively impaired participant has fewer and shorter visible distal cerebral arteries due to a longer blood transition time.

### 3.3 Association Between Neurovascular MRI Measurements and RBANS Scores

As shown in Table 3 and Figure 4, whole-brain dCAF index had a significant correlation with the RBANS Total score (*r*= 0.383, *p*= 0.028), and also with the immediate memory and attention index scores (*r*= 0.358, *p*= 0.041 and *r*= 0.384, *p*= 0.027, respectively). In a post hoc analysis adjusting for age and sex, whole-brain dCAF index still had a significant correlation with the RBANS Attention Index score (*r*= 0.380, *p*= 0.032), and a near significant correlation with the RBANS Total score (*r*= 0.347, *p*= 0.052).

**Figure 4.**
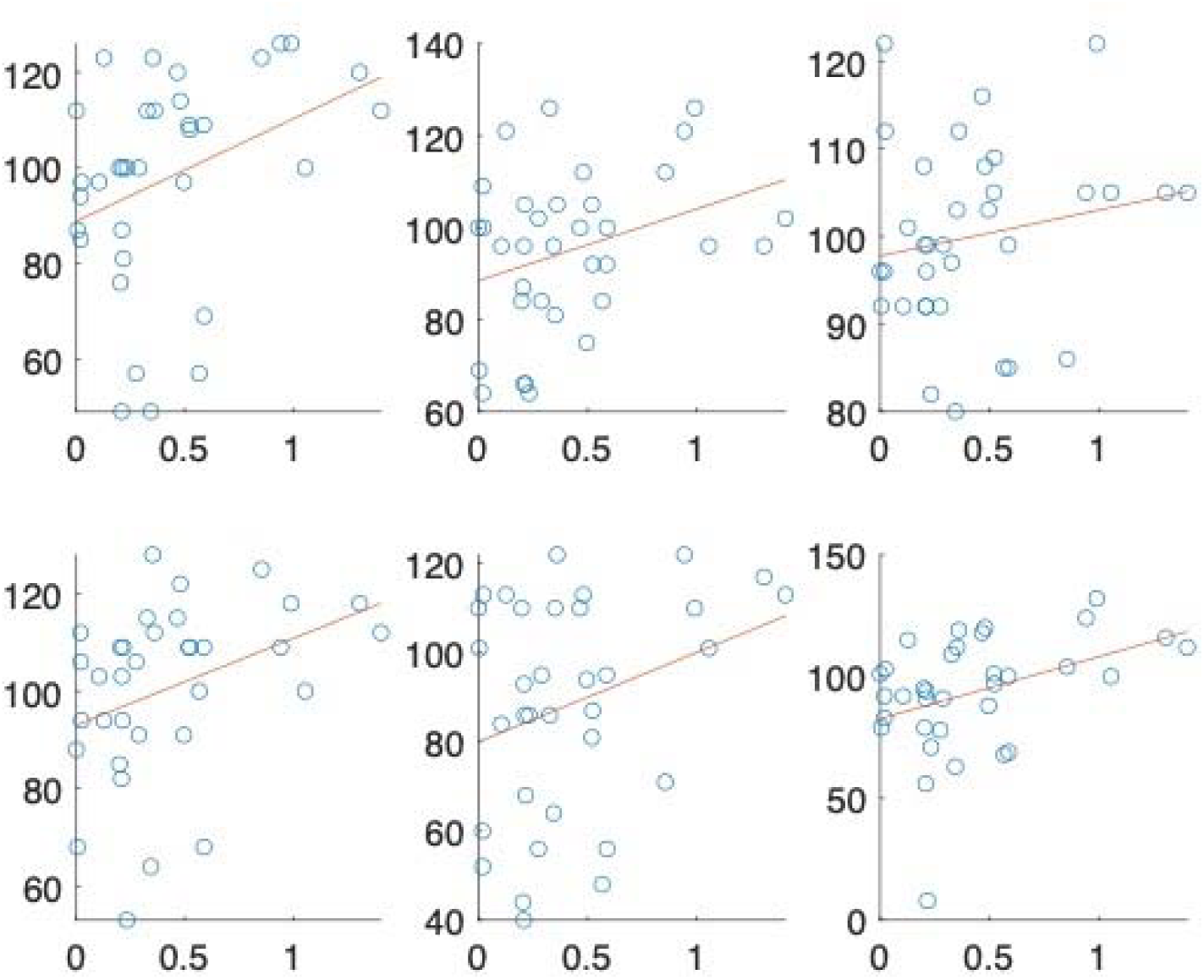
Scatter plots of whole-brain distal cerebral distal arterial flow (dCAF) index with domain-specific and total RBANS scores (N=35). The Pearson r and p values are marked on each plot (without adjusting for age and gender). There are significant associations between whole-brain dCAF index and immediate memory, attention and total RBANS scores.

**Table 3.** Partial Pearson correlations of whole-brain neurovascular MRI measurements with domain-specific and total RBANS scores.

| Neurovascular MRI Measurements | Model | Immediate Memory |  | Visuospatial Constructional |  | Language |  | Attention |  | Delayed Memory |  | Total RBANS Score |  |
| --- | --- | --- | --- | --- | --- | --- | --- | --- | --- | --- | --- | --- | --- |
|  |  | *r* | *p* | *r* | *p* | *r* | *p* | *r* | *p* | *r* | *p* | *r* | *p* |
| Distal Cerebral Arterial Flow (dCAF) index | 1 | 0.358 | **0.041\*** | 0.289 | 0.102 | 0.282 | 0.112 | 0.384 | **0.027\*** | 0.263 | 0.139 | 0.383 | **0.028\*** |
| (N=34) | 2 | 0.333 | 0.063 | 0.304 | 0.091 | 0.175 | 0.338 | 0.380 | **0.032\*** | 0.268 | 0.138 | 0.347 | 0.052 |
| Gray Matter CBF | 1 | 0.086 | 0.635 | 0.268 | 0.132 | 0.057 | 0.753 | 0.226 | 0.206 | -0.008 | 0.965 | 0.147 | 0.415 |
| (N=34) | 2 | 0.155 | 0.397 | 0.312 | 0.082 | 0.003 | 0.989 | 0.256 | 0.157 | 0.040 | 0.827 | 0.217 | 0.232 |
| Periventricular WMH (pWMH) Volume | 1 | -0.278 | 0.117 | -0.254 | 0.154 | -0.161 | 0.370 | -0.245 | 0.169 | -0.135 | 0.453 | -0.247 | 0.166 |
| (N=34) | 2 | -0.255 | 0.158 | -0.220 | 0.226 | -0.173 | 0.343 | -0.275 | 0.128 | -0.124 | 0.497 | -0.220 | 0.226 |
| Deep WMH (dWMH) Volume | 1 | -0.092 | 0.611 | -0.249 | 0.162 | -0.081 | 0.653 | -0.286 | 0.106 | 0.087 | 0.629 | -0.076 | 0.674 |
| (N=34) | 2 | -0.081 | 0.661 | -0.222 | 0.222 | -0.091 | 0.620 | -0.297 | 0.099 | 0.097 | 0.596 | -0.062 | 0.736 |
| Total WMH Volume | 1 | -0.230 | 0.197 | -0.261 | 0.143 | -0.142 | 0.432 | -0.265 | 0.137 | -0.072 | 0.689 | -0.204 | 0.255 |
| (N=34) | 2 | -0.209 | 0.251 | -0.228 | 0.210 | -0.153 | 0.403 | -0.290 | 0.108 | -0.060 | 0.743 | -0.179 | 0.327 |

\* Significant correlation ( $p < 0.05$ ); Model 1: without age and gender adjustment; Model 2 (post hoc analysis): with age and gender adjustment.

No significant correlations were found between gray matter CBF, pWMH, dWMH, or total WMH volumes and any RBANS scores (Table 3).

## 4 Discussion

In this study, we evaluated neurovascular MRI markers on an aging population and examined group differences between cognitively unimpaired and impaired (MCI or AD) groups. There were significant differences in the distal cerebral arterial flow (dCAF) index between the two groups, but not in gray matter CBF or WMH volumes. Further, the dCAF index was significantly associated with RBANS Immediate Memory Index, Attention Index, and Total scores.

### 4.1 Distal Cerebral Arterial Flow (dCAF) Index Measured From SNAP MRI

The total distal artery length visible in SNAP MRI has been shown to be associated with cognitive function in carotid atherosclerosis patients[18–20], with adjustment for brain size. In this study, we further define the total distal artery length normalized by intracranial volume as the distal cerebral arterial flow (dCAF) index, and employed it to assess cerebrovascular flow condition of older participants, not limited to vascular diseases, and obtained consistent results with previous studies: the dCAF index is significantly associated with the cognitive function (RBANS score), in the whole brain as well as in each arterial territory (except left PCA). This finding indicates that SNAP-based dCAF measurements may reflect the general cerebrovascular flow condition, not limited to the flow condition in atherosclerosis patients.

No significant correlation was found between age and dCAF measurement. This may be due to the limited range in age evaluated in the current study. And there is no significant difference in dCAF index between males and females. Despite no significant age or sex dependency for the dCAF index in this study population, we examined the correlation between dCAF index and RBANS scores with adjustment for age and sex as a post hoc analysis. After the adjustment, the correlation with RBANS Attention Index score remained significant while the correlation with RBANS Total score was nearly significant.

While the significant correlation between dCAF and RBANS Attention Index score found in this study appears consistent with clinical diagnoses, that is, early symptoms of vascular cognitive impairment can include difficulty paying attention[42], the finding is very preliminary due to a limited sample size. Furthermore, there can be an overlap between different sources of cognitive impairment in the same participant, for example, mixed vascular dementia and AD symptoms.

### 4.2 Gray matter CBF and WMH volume measurements

In addition to the newly-developed dCAF index, We included two established neurovascular MRI markers, gray matter CBF and WMH (total, pWMH and dWMH) volumes. In contrast to dCAF index, gray matter CBF neither demonstrated a significant between-group difference, nor showed a significant correlation with cognitive factors.

Interestingly, there is a significant correlation between dCAF index and gray matter CBF, in the whole brain (*r*= 0.342, *p*= 0.048) as well as in the left MCA territory (*r*= 0.428, *p*= 0.012), as shown in Figure S2. These findings suggest that although both are cerebral blood flow-related measurements, dCAF may present changes more sensitive to cognitive function, compared with CBF, which represents microvascular capillary flow.

The associations between WMH volumes and RBANS scores were unsignificant in this study, possibly because due to the limited sample size (*N*=34 after excluding missing values). The pWMH volume was found to be more associated with RBANS scores than total and dWMH volumes, which is consistent with the literature[35], since periventricular WMH may represent short brain connection disruptions that contribute more to cognitive function than deep WMH.

### 4.3 Distinct associations of neurovascular measurements with cognition outcome in the cognitively unimpaired vs. impaired participants

As a post-hoc analysis, we examined the Pearson correlation between each neurovascular measurement and the RBANS scores, separately for the cognitively unimpaired and impaired groups. Surprisingly, all measurements show an opposite relationship with RBANS score in the two groups, especially dCAF index and gray matter CBF (Figure S3). Though not significant, dCAF index and gray matter CBF have a positive correlation with RBANS score in the cognitively unimpaired group (*N*=22), and a negative correlation with RBANS score in the cognitively impaired group (*N*=13 and 12 for dCAF and CBF, respectively). The negative relationship between dCAF/CBF and cognitive function in the cognitively impaired group is counterintuitive and has not been reported before. One possible explanation is that there may exist a compensatory mechanism in MCI or AD subjects that triggers an elevation of the blood velocity, though further investigation is needed by including a larger sample size in both groups.

### 4.4 Limitations

One major limitation is that no neurovascular disease information (for example, carotid or intracranial atherosclerosis diseases) was collected for the participants, which were primarily enrolled in a larger neuroimaging study focused on the aging population.

Another limitation is that the number of participants is relatively small, and the MCI and AD participants were categorized in one group. Those limitations can be addressed in future studies by including vascular MRI measures (for example, MR angiography and vessel wall MRI), and by enrolling more participants and separating the MCI and AD participants in two different groups.

## 5 Conclusion

In conclusion, we evaluated MRI-based neurovascular measurements in an aging population, including a newly developed distal cerebral arterial flow (dCAF) index measured from the SNAP MRI technique. Statistical analyses suggested the dCAF index was significantly different between the cognitvely unimpaired and impaired participants, and it was significantly associated with cognitive function. This finding suggests SNAP-based dCAF index can potentially serve as an additional quantitative neurovascular marker for evaluating cognitive impairment and dementia.

## Acknowledgements

This work was supported by NIH K01-AG075166, NIH K01-AG073578, NIH R01-HL103609, NIH R01-NS127317, 2023 ADRC-USU Research Catalyst, and 2023 Center on Aging Seed Grant at University of Utah.

---

**Figure S1.**
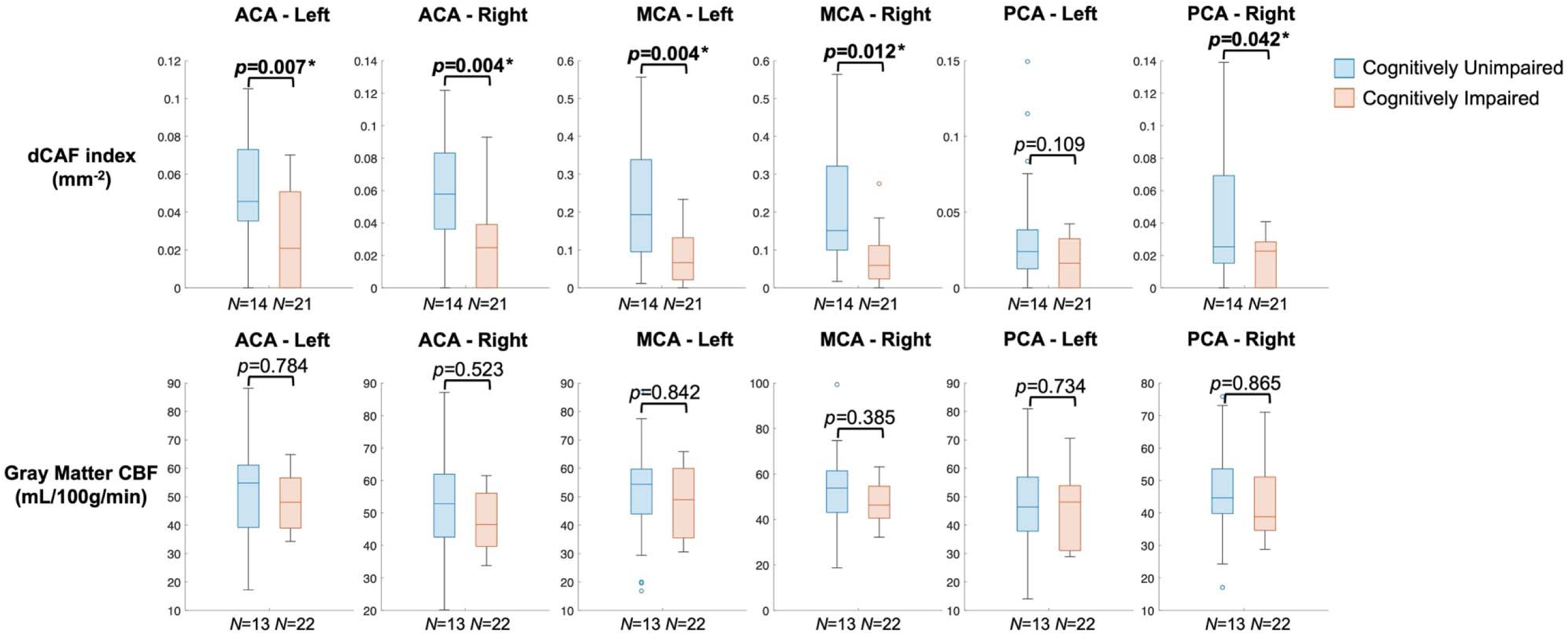
Comparison of territorial dCAF index and gray matter CBF between cognitively unimpaired and impaired groups. Shown is the box plot of each measurement displaying the median, upper and lower quartiles. There is a significant difference in dCAF index between the two groups for all territories except left PCA, while there is no significant difference in the territorial gray matter CBF between the two groups.

**Figure S2.**
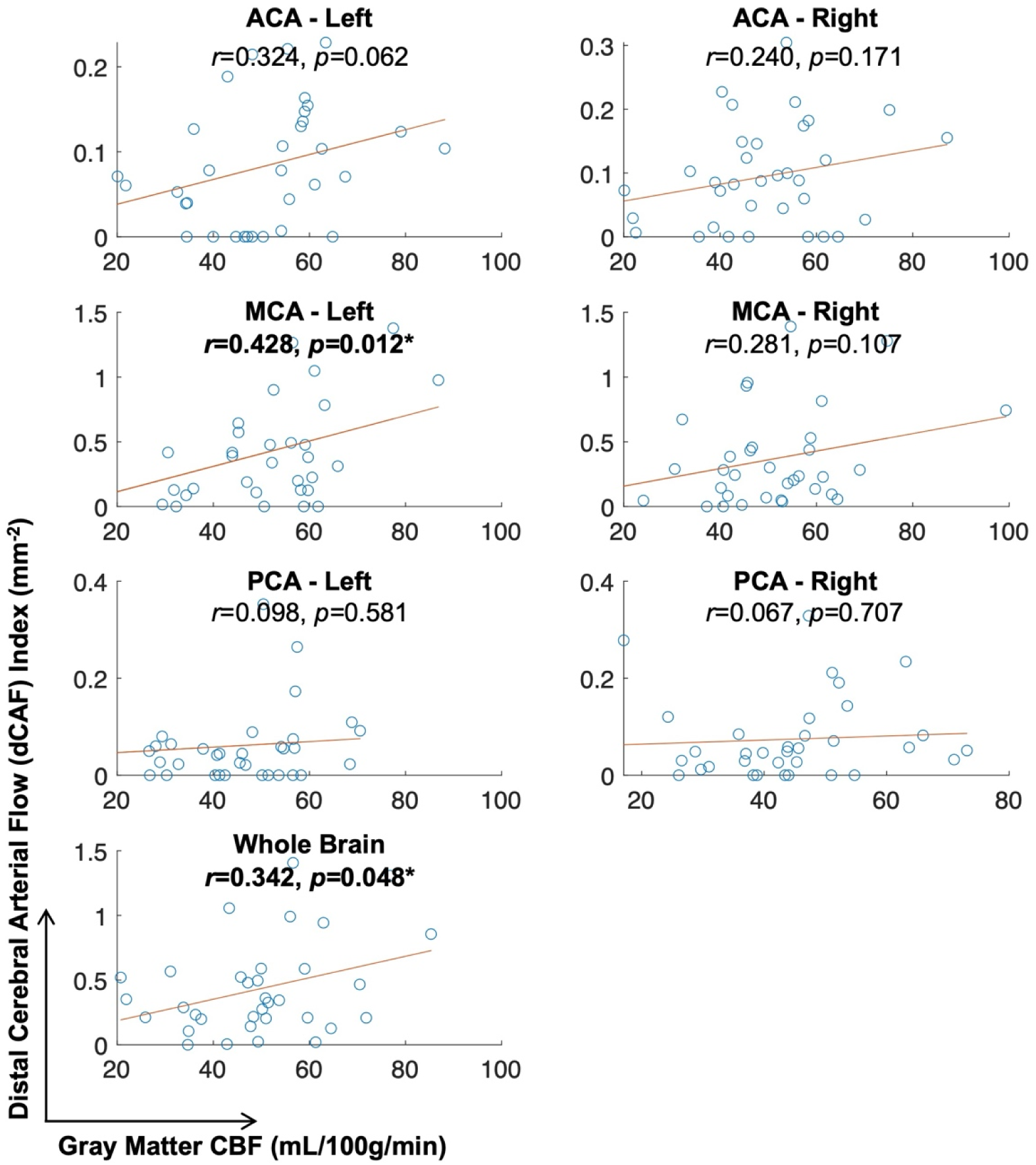
Results of Pearson correlation between distal cerebral distal arterial flow (dCAF) index and the gray matter CBF, in the whole brain as well as in each arterial territory (N=34). Significant associations were found between the two measurements in the whole brain and in the left MCA territory.

**Figure S3.**
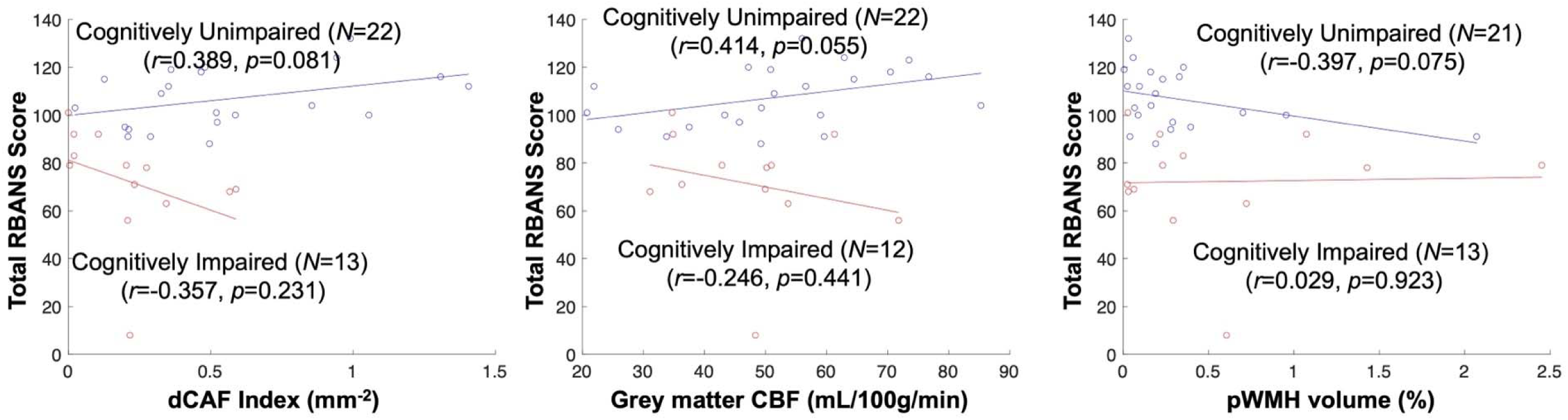
Group-specific Pearson correlation results between each whole-brain neurovascular MRI measurement and the total RBANS scores. Though not significant, there is a positive correlation between all MRI measurements and total RBANS scores in the cognitively unimpaired group, and a negative correlation in the cognitively impaired group.

